# The Nematode Antimicrobial Peptidome: a novel opportunity for parasite control?

**DOI:** 10.1101/2022.09.26.507570

**Authors:** Allister Irvine, Sharon A. Huws, Louise E. Atkinson, Angela Mousley

**Affiliations:** Microbes & Pathogen Biology, The Institute for Global Food Security, School of Biological Sciences, Queen’s University Belfast, 19 Chlorine Gardens, Belfast BT9 5DL, United Kingdom

**Keywords:** Antimicrobial peptide, AMP, nematode, parasite, bioinformatics, *in-silico*

## Abstract

Antimicrobial Peptides (AMPs) are key constituents of the invertebrate innate immune system where they provide critical protection against microbial threat. Knowledge of AMP complements within phylum Nematoda is limited however nematodes adopt diverse life strategies and frequently reside in microbe-rich environments such that they are likely possess broad AMP profiles with bioactivity against a range of microbiota. Indeed, parasitic nematode AMPs likely have roles in defence against invading pathogens and modulation of the host microbiome.

In this study the distribution and abundance of AMP-encoding genes were examined in 134 nematode genomes providing the most comprehensive profile of AMPs within phylum Nematoda. We reveal that phylum Nematoda is AMP-rich and -diverse, where 5887 genes encode AMPs. Genome and transcriptome analyses broadly reveal: (i) AMP family profiles that are influenced by nematode lifestyle where free-living nematodes appear to have an expansion of AMPs relative to parasitic species; (ii) major differences in the AMP profiles between nematode clades where Clade 9/V and 10/IV species possess expanded AMP repertoires; (iii) AMP families with highly restricted profiles (e.g. Cecropins and Diapausins) and others [e.g. Nemapores and Glycine Rich Secreted Peptides (GRSPs)] which are more widely distributed; (iv) complexity in the distribution and abundance of Defensin subfamily members; and (v) expression of AMPs in key nematode life stages.

These data indicate that phylum Nematoda has a diverse array of AMPs and underscores the need to interrogate AMP function to unravel their importance to nematode biology and host-worm-microbiome interactions. Enhanced understanding of the Nematode Antimicrobial Peptidome will inform drug discovery pipelines for pathogen control.

## 1. Introduction

Antimicrobial Peptides (AMPs) are natural immune effectors present in all classes of life (Wang, 2017). Invertebrate AMPs form the first line of defence against invading pathogens including bacteria, fungi and viruses by exerting broad-spectrum antimicrobial activities. AMPs also stimulate invertebrate immune regulation, enhancing the endogenous immune response to protect against microbial threat (van der Does et al., 2019). Whilst AMPs have gained significant attention as novel, resistance-breaking, antimicrobial agents in vertebrate medicine (Mwangi et al., 2019), their key roles in invertebrate innate immunity highlight an unexplored opportunity for the exploitation of AMPs as novel targets for the control of invertebrate pathogens, including nematode parasites. Current understanding of invertebrate AMP function is primarily derived from arthropods (Gomez et al., 2017) where they have been shown to be critical in defense against pathogens (Du et al., 2019; Zhang et al., 2021; Feng et al., 2022), however knowledge of the diversity, role and importance of AMPs, including in nematode parasites, is limited. Harnessing nematode-derived AMPs for parasite control demands a comprehensive understanding of AMP biology which is reliant on the characterisation of AMP diversity in key nematode pathogens.

Parasitic nematodes frequently reside in hazardous host environments such as the microbe-rich intestine where they survive in close association with a broad range of microorganisms such that gastrointestinal nematodes are likely to require a raft of endogenous AMPs for survival within the host. Knowledge of AMPs in nematodes has been derived from research using the model organism *Caenorhabditis elegans* and the gastrointestinal pig parasite *Ascaris suum*, and more recently broader *in silico* studies, where a number of AMP families have been identified (Cecropins, Defensins, Nemapores, Diapausins, Glycine Rich Secreted Peptides (GRSPs) (Lee et al., 1989; Kato and Komatsu, 1996; Banyai and Patthy, 1998; Andersson et al., 2003; Couillault et al., 2004; Dieterich et al., 2008; Tarr, 2012a; Ying et al., 2016; Gu et al., 2018). The ability to uncover AMPs in nematodes has been significantly enhanced by the availability of helminth genome and transcriptome datasets (Doyle, in press). Indeed, prior to this study, the most complete profile of AMPs in nematodes was conducted by Tarr (2012a), where draft genomes and Expressed Sequence Tag (EST) datasets were examined in 58 nematode species for a sub-set of AMP families (Cecropin, Defensin, Nemapore). A constraint, noted by Tarr (2012a), was the lack of publicly available nematode ‘omics data which prevented the comprehensive characterisation of nematode AMP families. Recent expansions and quality improvements in nematode ‘omics data present a timely opportunity for the characterisation of pan-phylum nematode AMP diversity that will provide a springboard to functional biology and therapeutic exploitation.

This study examines 134 nematode genomes, representing 109 species, to profile five AMP-encoding gene families (Cecropin, Defensin, Diapausin, Nemapore, GRSP) using a homology-directed bioinformatics approach. To our knowledge, the data presented here represent the most comprehensive profile of nematode AMPs to date and signify a critical advance in our understanding of the nematode AMP repertoire. These data expose nematodes as a rich source of AMP diversity that displays both conservation and variation within and between nematode clades, lifestyles and life stages likely reflecting biologically relevant trends.

These data reveal AMP diversity within phylum Nematoda and provide a novel opportunity to probe AMP function in therapeutically relevant parasites to underpin novel nematode control approaches. In addition, this study has provided a novel library of nematode-derived natural antimicrobials which may exhibit desirable characteristics for future antimicrobial discovery.

## 2. Materials and Methods

### 2.1. Identification of putative AMP-encoding genes in phylum Nematoda

A summary of the homology-directed methods in this study have been provided graphically (see Figure 1A). Genes encoding putative AMPs representing five families (Cecropin, Diapausin, Defensin, Nemapore and GRSP) were identified based on homology to previously identified nematode AMPs using Hidden Markov Model (HMM) and Basic Local Alignment Search Tool (BLAST) searches. HMM profiles were built using HMMER v3.2.1 (www.hmmer.org) for each AMP family and, where appropriate, for individual subfamilies. Previously identified AMPs (see Supplementary Table S1) were aligned using CLUSTAL Omega with default settings (Madeira et al., 2019), followed by *hmmbuild* (with default settings) to construct specific AMP family HMM profiles. Where required, alignments were manually adjusted to align AMP motifs. A database of nematode predicted proteins was collated from the predicted protein datasets derived from 134 nematode genomes (Wormbase Parasite V14; Supplementary Table S2; Bolt et al., 2018; Harris et al., 2020). Constructed HMM models were subsequently searched against the concatenated nematode predicted protein database using the *hmmsearch* command. The results of each HMM search were cross-checked to ensure that the query sequences used to build the model scored highly. This confirmed that the HMM profile successfully identified the query sequences and was specific for each AMP family. For all AMP families, except the GRSPs, search hits with an E value <0.01 were manually assessed for high peptide similarity and conserved AMP family motifs. Where a sequence appeared incomplete, as a result of an assembly or annotation error (i.e., missing an exon or incorrectly predicted as a longer protein), the output was retained to provide a comprehensive profile of putative AMP-encoding genes. For the Cecropin family, new hits were included based only on homology to the query sequences, whereas for the Defensin, Diapausin and Nemapore families hits were included based on the presence of specific conserved cysteine motifs. As the GRSPs are characterised by repetitive glycine-rich repeats and lack other conserved motifs, individual subfamily HMMs were constructed based on the 17 GRSP subfamilies identified in *C. elegans* and *Caenorhabditis briggsae* GRSPs (Ying et al., 2016). Due to the highly repetitive glycine-rich nature of the GRSPs, all hits from these HMMs up to an E value of 10 were assessed and were confirmed based on GRSP criteria established by Ying et al. (2016). As a result, putative GRSP hits were only considered to be authentic if they were ≤200 amino acids, contained a putative signal peptide sequence, and possessed mature peptide glycine content >17%. Despite the construction of 17 subfamily HMM profiles for the GRSPs, results overlapped making classification of putative GRSPs according to the pre-defined *Caenorhabditis* GRSP subfamilies difficult. Consequently, GRSPs hits were not classified into discrete subfamilies in this study.

**Figure 1:**
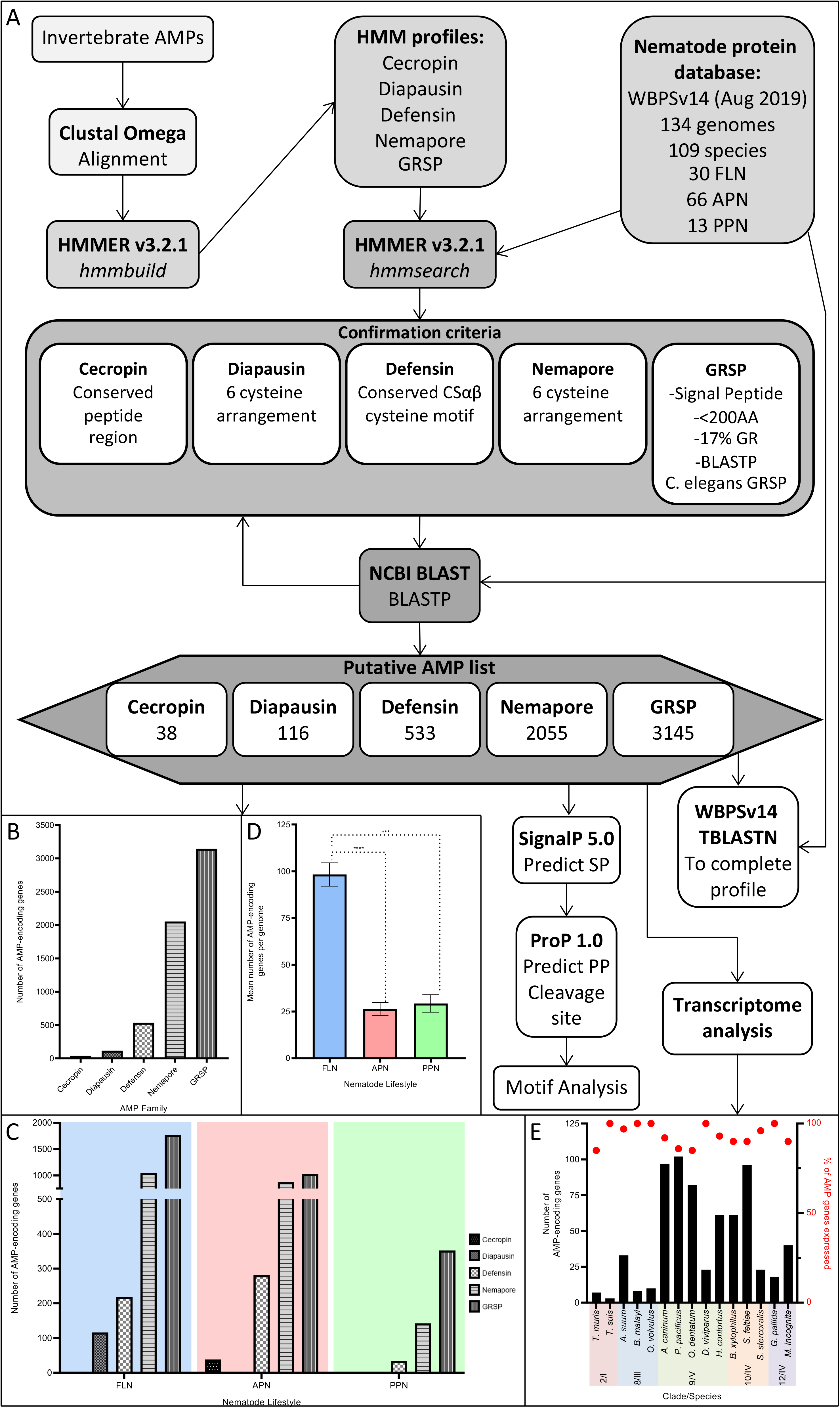
An *in silico* AMP discovery pipeline reveals that phylum Nematoda is a rich source of AMPs. A) Summary of homology-based methods for putative AMP identification. B) Number of AMP-encoding genes identified for each nematode AMP family. C) Mean number of AMP genes per genome for each AMP family across nematode lifestyles. D) Total number of AMP-encoding genes across each nematode lifestyle. E) Summary of AMP-gene expression (> 2TPM) relative to the number of AMP-encoding genes for key nematode species. Error bars indicate standard error of the mean. *** P<0.001; **** P<0.0001. AA, Amino Acid; AMP, Antimicrobial Peptide; APN, Animal Parasitic Nematode; BLAST, Basic Local Alignment Search Tool; BLASTP, Protein BLAST; CSαβ, Cysteine-Stabilised Alpha Beta peptide superfamily; FLN, Free-Living Nematode; GR, Glycine Rich; GRSP, Glycine Rich Secreted Peptide; HMM, Hidden Markov Model; PP; Propeptide; PPN, Plant Parasitic Nematode; SP, Signal Peptide; TBLASTN, Translated nucleotide BLAST; WBPSv14, WormBase Parasite Version 14

Post HMM analyses, all HMM-derived putative AMPs were employed as BLASTp query sequences in the NCBI command line BLAST application (version 2.9.0, April 2019) against the concatenated nematode predicted protein database constructed previously. BLASTp returns were confirmed using the same criteria as above for each AMP family and new confirmed hits were repeatedly used as BLASTp queries until no new results were retrieved. Due to the high number of putative GRSP hits returned by HMM and BLASTp searches and the potential for false-positive returns, all putative GRSP hits were subjected to an additional BLASTp search against the *C. elegans* protein database on the Wormbase Parasite BLAST server (with the low complexity filter off; Version 14); where a GRSP identified previously by Ying et al. (2016) was not returned within the top five hits of the additional BLASTp search (sorted by E value) the original putative GRSP hit was removed from further analyses.

Putative AMP prepropeptides were analysed for the presence of signal peptide sequences and propeptide cleavage sites using SignalP 5.0 (standard Sec/SPI settings, Eukarya organism; Armenteros et al., 2019) and ProP 1.0 (default furin-specific prediction; Duckert et al., 2004) respectively.

The Defensin family was subsequently classified into subfamilies (Mollusc/Nematode Defensin, Traditional Insect Defensin, Macin and Drosomycin) based on number of cysteines and their arrangement according to the cysteine reference array established previously (Tarr, 2016).

To reduce the possibility of false-negative returns post BLASTp, tBLASTn searches using query sequences from closely related species with a positive hit were employed (via Wormbase Parasite V14 BLAST server); this approach confirmed null returns and checked for the presence of unannotated sequences. Where an unannotated putative AMP was identified, the nematode AMP profile was updated to reflect this however unannotated genes could not be included in relative abundance or transcriptome analyses. Due to high sequence variability and gene expansions in specific genera it was not possible to accurately assign AMP genes as orthologs.

### 2.2. Transcriptome analyses of AMP-encoding genes

Publicly available life-stage specific RNAseq datasets for 15 nematode species, representing a range of nematode clades and lifestyles, were retrieved from Wormbase Parasite v14 (see Supplementary Table S2). A threshold of ≥2 Transcripts Per Million (TPM) was employed as a cut-off for expression (Wagner et al., 2013). For heatmap construction TPM values were Log2 transformed and converted to Z values using Heatmapper (www.heatmapper.ca/; Babicki et al., 2016). For Log transformation and heatmap visualisation TPM 0 was reported as TPM 0.01. Heatmaps were scaled by row and genes were clustered by row using the Average Linkage method with Euclidean Distance Measurement method.

### 2.3. Statistical analyses

All graphs and statistical analyses were produced using GraphPad Prism 9 (www.graphpad.com/). Data were tested for normality using the Kolmogorov-Smirnov test (Massey Jr, 1951). Non-normally distributed data were tested for significance using Krustal Wallis Test with Dunn’s multiple comparisons test (Dunn, 1961).

## 3. Results and Discussion

### 3.1. Phylum Nematoda is a rich source of antimicrobial peptides

To investigate AMP abundance and diversity across phylum Nematoda, five known nematode AMP families (Cecropin, Diapausin, Defensin, Nemapore and GRSP (Tarr, 2012b; Zhu and Gao, 2014; Ying et al., 2016; Gu et al., 2018) were profiled in 109 nematode species (134 genomes; representing 7 clades, 30 free-living nematode (FLN), 66 animal parasitic nematode (APN) and 13 plant parasitic nematode (PPN) species; see Supplementary Table S2). Our analyses identified 5887 putative AMP-encoding genes, of which >90% are novel, representing all known nematode AMP families [38 (0.65%) Cecropin-encoding genes, 116 (1.97%) Diapausin-encoding genes, 533 (9.05%) Defensin-encoding genes, 2055 (34.91%) Nemapore-encoding genes and 3145 (53.42%) GRSP-encoding genes; see Figure 1B and Supplementary Table S3)]. Whilst the relative abundance and profile of AMP-encoding genes vary across the phylum (see Figure 2) the representation of AMPs in every nematode species examined underscores the importance of AMPs to nematode biology. These data provide the most comprehensive profile of AMPs across the phylum Nematoda and demonstrate that nematodes are AMP-rich.

**Figure 2.**
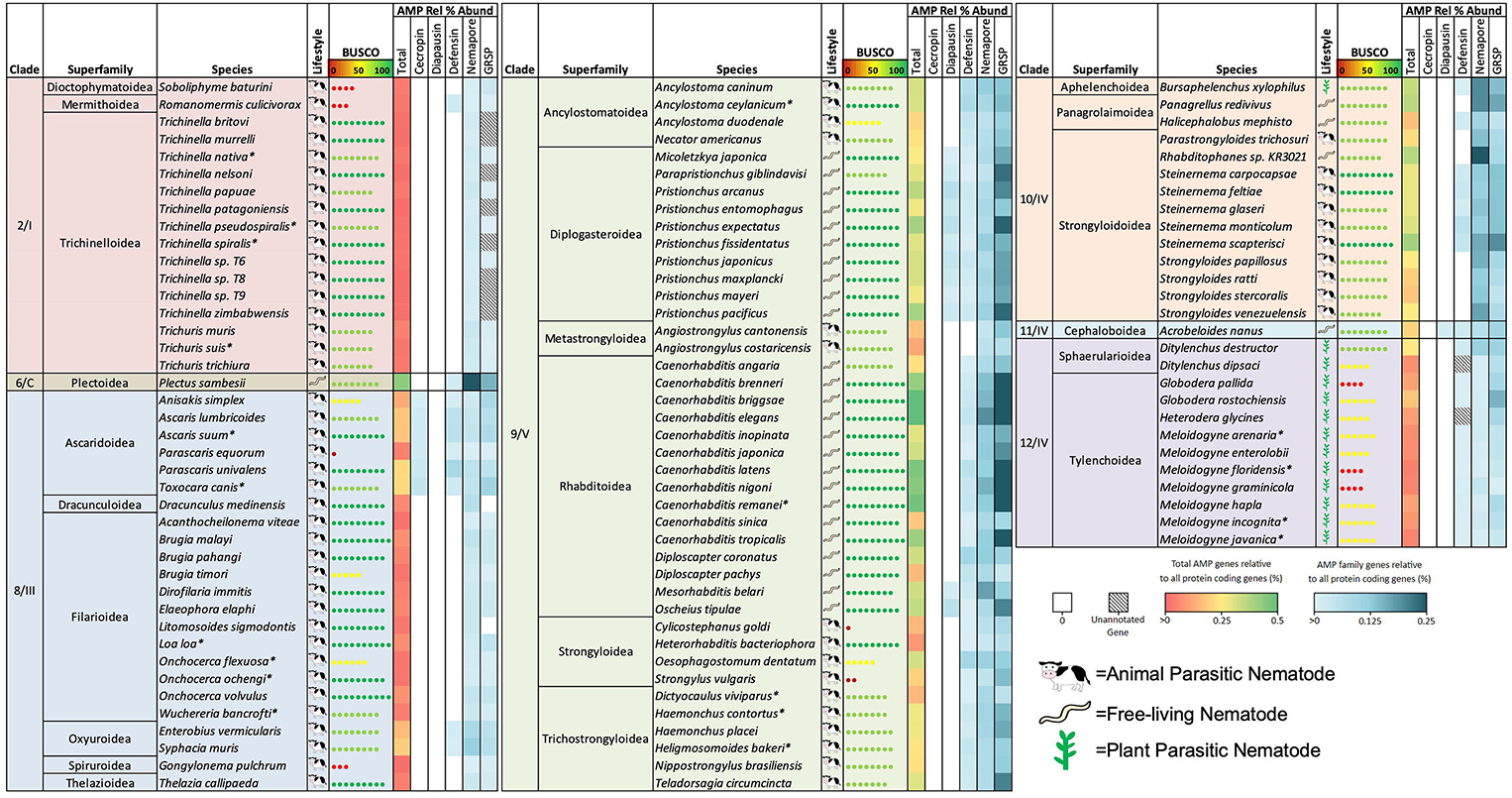
Relative abundance of AMP genes reveal AMP profile diversity across nematode clades and lifestyles. Nematode species (109 nematode species; 134 genomes) are organised by clade (Blaxter et al., 1998, van Megan et al 2009) and superfamily designation (NCBI taxonomy browser; https://www.ncbi.nlm.nih.gov/taxonomy). Relative abundance was calculated as a percentage of AMP genes relative to total protein coding genes per genome. BUSCO complete scores were obtained from Wormbase Parasite V14 (https://parasite.wormbase.org/index.html; Bolt et al 2018). Asterisk indicates multiple genome assemblies were used.

### 3.2. AMP profile, abundance, and family diversity vary across nematode lifestyles

Our analyses reveal that AMP-encoding gene family profiles vary across nematode lifestyles (see Figure 1C). FLN and APN species display the greatest AMP family diversity (Diapausin, Defensin, Nemapore and GRSP families represented in FLNs; Cecropin, Defensin, Nemapore and GRSP families represented in APNs), however PPNs lack both Diapausin and Cecropin families.

FLNs also possess the greatest number of AMP-encoding genes (98.31 ± 6.23 genes per genome) relative to both APNs (26.37 ±3.53 genes per genome; p<0.0001) and PPNs (29.33±4.67 genes per genome; p= 0.0001) (see Figure 1D). The higher abundance of AMP-encoding genes in FLNs is not surprising given the diversity in the terrestrial and marine environments that they inhabit, and in the heterogeneity in microbes that they are likely to encounter. Parasitic nematodes, in contrast, live endogenously within animals and plants and are likely to benefit from host immunity which may somewhat protect against microbial threat reducing the reliance on AMPs. Parasitic nematodes, however, also likely possess an AMP arsenal which is more specialised to host-dwelling microbes and pathogens.

### 3.3. Clade 9/V and 10/IV nematodes are AMP-rich

There was considerable variability in the distribution and abundance of AMP families across nematode clades (see Figure 2). For clade comparisons Clades 6/C and 11/IV were excluded as there is only a single species with a publicly available genome representing each of these clades [*Plectus sambesii* (Clade 6/C); *Acrobeloides nanus* (Clade 11/IV)]. Both of these species are free-living and display high AMP diversity (see Figure 2); expansion of genome datasets is required to determine whether the diverse AMP profiles noted here are conserved traits for Clade 6/C and Clade 11/IV species.

Clade 9/V species displayed the highest AMP diversity across the phylum (80.77 ±5.736 AMP-encoding genes per genome). To account for potential differences in comparisons between genomes with different gene numbers, the number of AMP genes was calculated relative to the total number of protein coding genes (PCGs) for each genome (see Figure 2 and Supplementary Table S4). By this measure, Clade 9/V displayed the highest relative abundance of AMP genes (average 0.31 ±0.02% per genome). Within Clade 9/V, there were differences between the relative abundance of AMP genes across species where *C. elegans* had the highest AMP complement (0.82% of total number of PCGs). In contrast, the lowest relative abundance of AMP genes was noted for the entomopathogenic nematode (EPN), *Heterohabditis bacteriophora*, where only 0.09% of PCGs encode AMP genes. The high abundance of AMP genes in *C. elegans* was unsurprising given that this species has provided the blueprint for nematode AMP research. Indeed, most of the query sequences used to identify AMP genes in this study originated from *C. elegans*. The reduced relative abundance of AMP genes in *H. bacteriophora* may be related to its symbiotic relationship with gram negative *Photorhabdus* bacteria (Parihar et al., 2022). *Photorhabdus* spp. are key to the lifecycle of *H. bacteriophora* and produce a range of toxins which kill the arthropod host (Clarke, 2020). Recently, an antibiotic with potent activity against gram negative bacteria was discovered in *Photorhabdus* (Imai et al., 2019). It is therefore possible that *H. bacteriophora* benefits from AMPs produced by the bacterial symbiont reflecting the lower abundance of endogenous nematode AMPs in this species.

Beyond the AMP-rich *Caenorhabditis* genera, other genera within Clade 9/V also appear to display enriched AMP profiles. For example, free-living *Pristionchus* species have an enriched AMP profile with all but one species having >100 AMP genes (0.33 ±0.02% of all PCGs). *Pristionchus* feed on rotting plant matter and share mostly phoretic, and occasionally necromenic, relationships with arthropods (Félix et al., 2018). It may be the case that *Pristionchus* nematodes require an enriched arsenal of AMPs, to combat the microbe-rich environments they inhabit.

The Ancylostomatoidea and Trichostrongyloidea superfamilies, which include mammalian parasites, also have a higher AMP abundance than the other parasitic superfamilies, e.g. the Metastrongyloidea and Strongyloidea, in this clade. Many of these parasitic species that display higher AMP abundance, including *Necator americanus, Teladorsagia circumcincta* and *Haemonchus contortus*, are known to alter the host gut microbiome during infection (El-Ashram et al., 2017; Cortés et al., 2020; Jenkins et al., 2021). Whilst it is currently unclear whether AMPs from these species are secreted and/or excreted into the host environment to contribute to changes in the microbiome, nematode-derived AMPs, such as *A. suum* Cecropin *P1*, have been isolated from the host gut (Lee et al., 1989; Andersson et al., 2003). Future research should aim to establish whether excreted/secreted nematode AMPs are active against host microbiota and contribute to changes in the host microbiome during infection.

Clade 10/IV is also AMP-rich (64.79 ±9.51 AMP-encoding genes per genome; 0.28 ±0.02% of PCGs in the Clade 10/IV species examined in this study encode AMPs). Despite the diversity in lifestyles, most Clade 10/IV species share similar numbers of AMPs. *Steinernema* spp. appear to have an elevated AMP abundance relative to other Strongyloidoidea superfamily members such as *Strongyloides* spp. Interestingly, like Clade 9/V *H. bacteriophora, Steinernema* spp. are also EPNs that display endosymbiotic relationships with gram negative bacteria. *Steinernema* spp. harbour *Xenorhabdus* bacteria which are critical for worm establishment and ultimately kill the insect host (Goodrich-Blair, 2007). Consistent with the relationship between *Photorhabdus* and *H. bacteriophora, Xenorhabdus* also produces antimicrobials which aid establishment and maintenance of the nematode parasites in the host (Dreyer et al., 2019). *Steinernema* spp. however do not appear to have the same reduction in AMP gene abundance as was observed in *H. bacteriophora*. At present, little is known about the role that worm-derived AMPs play in the lifecycle of EPNs. A recent excretory/secretory protein (ESP) analysis of *Steinernema feltiae* activated infective juveniles identified 266 proteins which were designated as ‘venom’ proteins involved in host infection (Chang et al., 2019); here we reveal that seven of these ESPs are Nemapores (L889_g29545, L889_g13905, L889_g31036, L889_g31037, L889_g17551, L889_g20297 and L889_g6015). Notably, these Nemapores were absent from the ESP of non-activated infective juveniles demonstrating that they likely play a role in *S. feltiae* infection and are actively secreted/excreted from the worm upon exposure to host tissue. The original study (Chang et al., 2019) did not identify these genes as Nemapores based on automatic annotation, demonstrating the value of the manual homology-directed approach used here. The presence of Nemapores in *S. feltiae* ESP also encourage more focussed peptidomic analyses of parasitic nematode ESP products to determine host/environment facing AMP profiles.

### 3.4. Clade 2/I, 8/III and 12/IV nematodes possess less diverse AMP profiles

Multiple nematode clades possess a limited AMP profile in comparison to Clade 9/V and 10/IV in terms of the number of AMP genes and the distribution of AMP families. Clade 2/I nematodes possess the most limited AMP profile of all the nematode clades profiled in this study (3.44 ±0.50 AMP-encoding genes per genome; 0.02 ±0.01% of PCGs encoding AMP genes). All species from this clade possess genes encoding Nemapores and GRSPs with only *Romanomermis culicivorax* possessing Defensin-encoding genes (see Figure 2). Clade 2/I nematodes are generally considered to be more basal and have more limited gene complements for other protein families including neuropeptides, antioxidant enzymes, G-protein-coupled receptors (GPCRs) and ATP-binding cassette transporters (ABC transporters) (McCoy et al., 2014; IHGC 2019; Xu et al., 2020; McKay et al., 2021). The reduced AMP arsenal observed in Clade 2/I may indicate that elevated AMP diversity in the Crown Clades (Clade 8-12) originated as a result of gene duplications and/or selective pressures that are not observed in the basal clades. It is interesting that many Clade 2/I nematodes share similar host niches with some of the parasitic species that have more diverse AMP profiles. Indeed, both *Trichinella* and *Trichuris* species spend much of their lifecycles in the microbe-rich gut environment of their hosts. The reduced number of AMP-encoding genes in Clade 2/I nematodes poses an intriguing question about how these nematodes combat microbial threats without the range of AMPs that other intestinal parasitic nematodes possess. It is possible that Clade 2/I species may possess divergent AMPs or larger antimicrobial proteins that cannot be identified using the homology-based pipeline employed in this study.

With the exception of the Ascaridoidea (31.38 ±4.14 AMP-encoding genes per genome; 0.17 ±0.02% of PCGs encoding AMPs) and Oxyuroidea (20.50± 0.50 AMP-encoding genes per genome; 0.17 ± 0.02% of PCGs encoding AMPs) superfamilies, Clade 8/III nematodes also broadly displayed a reduced AMP complement (6.95 ±0.01 AMP-encoding genes per genome; 0.06 ± 0.01% of PCGs encoding AMPs) in comparison to Clade 9/V species. This reduced AMP complement was particularly evident within the Filarioidea species which may suggest that there has been a loss of AMP genes from filarial nematodes. In line with the more basal nematodes in Clade 2/I, reduced profiles of metabolic and neuropeptide pathway components are also observed in filarial nematodes (IHGC 2019; McKay et al., 2021). Filarial nematodes typically reside in the circulatory system and subcutaneous tissues of the definitive host and may, therefore, be less reliant on AMPs due to the more sterile host niches they occupy. Additionally, many filarial nematodes including *Brugia malayi, Onchocerca volvulus* and *Wuchereria bancrofti* share well characterised endosymbiotic relationships with *Wolbachia* which are critical for key aspects of nematode biology (Taylor et al., 2005). It is possible that *Wolbachia* provides a level of protection for filarial parasites such that the nematodes require fewer endogenous AMPs. Alternatively, the reduced AMP profiles in Filarids could help shape a favourable environment for *Wolbachia* growth.

Clade 12/IV species, which include the PPNs, also possess a reduced AMP profile (27.47± 4.54 AMP-encoding genes per genome; 0.09± 0.01% of PCGs encoding AMPs) in comparison to Clade 9/V nematodes. Notably, the only PPN in this study that is not a Clade 12/IV species, *Bursaphelenchus xylophilus* (Clade 10/IV), possesses 61 AMP-encoding genes (0.34% relative abundance), exhibiting a profile much more in line with Clade 10/IV species. As noted for Clade 2/I nematodes there is also potential that there are currently undiscovered AMP families within Clade 12/IV species with diverse motifs that are not identifiable via the methods employed here. This highlights the need to expand AMP studies beyond *C. elegans* and *A. suum* to discover more evolutionarily distinct AMP families.

### 3.5 Cecropin genes are entirely restricted to Ascaridoidea species

The pan-phylum data presented here provides evidence to support the restriction of Cecropin-encoding genes to Ascarid species; these data also report the first identification of Cecropin-encoding genes from *Parascaris* spp and confirm the recent identification of Cecropins from *Anisakis* species (Rončević et al., 2022). The restriction of Cecropins to Ascaridoidea suggests Ascarid-specific roles that require further characterisation. As noted above, Cecropins were originally isolated from the porcine intestinal environment (Lee et al., 1989). Despite this, the effect of these peptides on host microbiota has yet to be characterised.

Cecropins have not undergone the same gene expansion observed in other AMP families (4.75 ± 0.88 genes per genome). The Ascarid Cecropins appear to be highly conserved suggesting that these peptides have similar antimicrobial activities (Figure 3). Indeed, chemically synthesised *A. suum* Cecropin P1-P4 have similar activity profiles against bacteria and fungi (Pillai et al., 2005). However, minor differences in amino acids can significantly change the antimicrobial activity of these peptides; for example, a single amino acid substitution of a Proline residue for Isoleucine residue at position 22 of *A. suum* Cecropin P1 significantly reduces antibacterial and membrane permeabilising activity (Gazit et al., 1996). Antimicrobial activity should, therefore, be investigated for Cecropin peptides from other Ascarid species.

**Figure 3:**
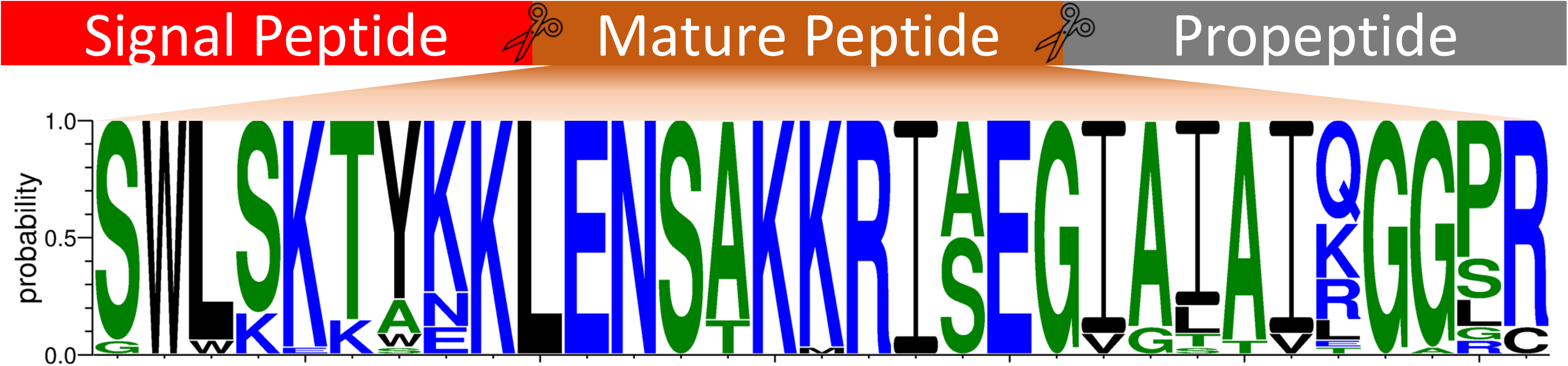
Ascaridoidea-restricted Cecropins display significant amino acid conservation. Weblogo generated using 34 cecropin peptide sequences (4 peptides excluded due to gene mispredictions; GS_20806, TCNE_0000065801, Tcan_02070, GS_06967) using WebLogo3 (https://weblogo.threeplusone.com/), coloured according to amino acid hydrophobicity (hydrophilic=blue, neutral=green, hydrophobic=black). Amino acid letter height reflects the probability of conservation at a specific location. Signal peptide, mature peptide and propeptide indicated.

### 3.6. Diapausin genes are restricted to specific free-living soil nematodes

Nematode Diapausin-encoding genes were originally identified from *Pristionchus pacificus* (Dieterich et al., 2008). The pan-phylum analyses performed here reveals that Diapausins are restricted to Clades 9/V and 11/IV in soil-dwelling FLNs, primarily the Clade 9/V Diplogasteroidea which includes *Pristionchus* spp. Beyond the Diplogasteroidea, two Rhabditoidea, *Mesorhabditis belari* and *Oscheius tipulae*, also encode Diapausins while the closely related *Caenorhabditis* genera do not. The absence of Diapausin-encoding genes in *C. elegans* is interesting given the similar environmental niche occupied by these species, highlighting the importance of AMP research beyond *C. elegans*. Diapausin-encoding genes were also identified in *Acrobeloides nanus*, the only Clade 11/IV species in this study; this is an intriguing observation from an evolutionary perspective.

It has been suggested that *P. pacificus* Diapausin-encoding genes have been acquired through horizontal gene transfer from beetles with which they share phoretic/necromenic associations (Rodelsperger and Sommer, 2011). This theory is supported by the presence of a Diapausin-like sequences in an insect iridovirus implicating the virus as the vector for gene transfer (Tanaka et al., 2003). Phylogenetic analyses did not provide any further insight into these relationships or gene evolution (data not shown) primarily due to poor resolution as a consequence of the rapid evolution of AMP-encoding genes.

In total 116 Diapausin-encoding genes were identified across phylum Nematoda with an average of 8.92 ±1.62 genes per genome. Diapausins possess a cysteine motif that consists of six cysteine residues which form three disulfide bonds (see Figure 4A). The position of these cysteines is conserved compared to arthropod Diapausins, suggesting conservation in Diapausin tertiary structure. Interestingly, some of the Diapausin-encoding genes identified in this study possessed two complete cysteine Diapausin motifs (e.g. ACRNAN_scaffold769.g15805; 12 cysteine residues). So far, Diapausin-encoding genes have only been reported to possess a single Diapausin cysteine motif and thus it is unclear whether the genes identified here contain multiple domains from errors in gene annotation or whether they represent a novel type of Diapausin-encoding gene, where multiple peptides could be cleaved from a single gene precursor. Beyond the nematode Diapausin cysteine motif there is significant amino acid variation that may result in different antimicrobial activity profiles between individual peptides. While Diapausins in beetles are potent antifungal agents which play a role in diapause (Tanaka et al., 2003), the role of Diapausins in nematodes has yet to be established.

**Figure 4:**
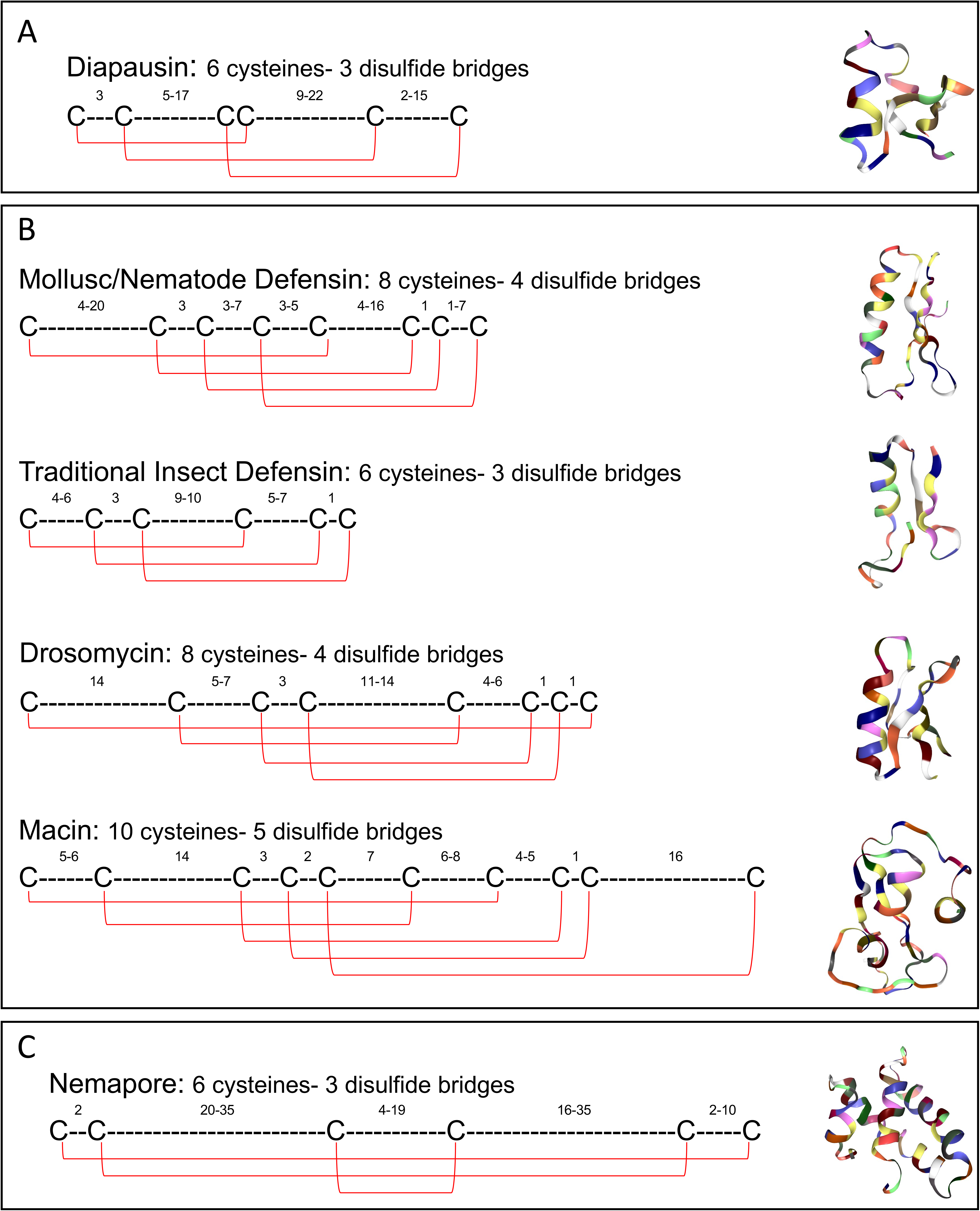
Cysteine organisation of the nematode cysteine-stabilised AMP families enables delineation of AMP subfamilies. A: The Diapausin cysteine organisation. Peptide family 3D structure example: *Gastrophysa atrocyanea* Diapause-specific peptide (RCSB: 2E2F). B: Organisation of cysteine residues in the CSαβ Superfamily members. Peptide family 3D structure examples: Mollusc/Nematode Defensin (*Ascaris suum* ASABF-α; RCSB:2D56), Traditional Insect Defensin (*Sarcophaga peregrina* Sapecin; RCSB: 1L4V), Drosomycin (*Drosophila melanogaster* Drosomycin; RCSB: 1MYN), Macin (*Hirudo medicinalis* Theromacin; RCSB:2LN8). C: The Nemapore cysteine organisation. Peptide family 3D structure example: *Caenorhabditis elegans* Caenopore-5 (RCSB:2JS9). ‘C’ indicates a Cysteine residue. ‘-’ indicates any other amino acid. Subscript numbers indicate the range in number of amino acids between each cysteine residue. Red lines indicate putative disulphide bonds between cysteine residues. Peptide family 3D structures obtained from Research Collaboratory for Structural Bioinformatics Protein Data Bank (RCSB PDB) (https://www.rcsb.org/; Berman et al., 2000).

### 3.7. Classification of nematode Defensins reveals complexity in subfamily distribution

Defensins are a superfamily of cysteine-stabilised alpha-beta (CSαβ) peptides which share a tertiary structure comprising an α-helix and two antiparallel β-sheets stabilised by three disulfide bonds (Zhu et al., 2005). Defensins can be classified into multiple groups of AMPs displaying differences in cysteine organisation. 533 Defensin-encoding genes (3.98 ±0.50 genes per genome) were identified in this study and were broadly distributed across phylum Nematoda, with representatives in every nematode clade (see Figure 2). Defensin-encoding genes are more restricted in Clades 2/I (only *R. culicivorax*) and 8/III (only Ascaridoidea and Oxyuroidea spp.), which is consistent with a previous study (Tarr, 2012a). In contrast, Defensin-encoding genes are more widespread in Clades 9/V, 10/IV and 12/IV (see Figure 2); note that two Defensin-encoding genes, previously identified from *P. trichosuri* EST datasets (Tarr, 2012a), were not identified here.

Defensin-encoding genes identified in this study were classified into four subfamilies based on the cysteine reference array reported previously (Tarr, 2016): Mollusc/Nematode Defensins (MN), Traditional Insect Defensins (TIDs), Macins and Drosomycins. The distribution of these subfamilies was variable across phylum Nematoda (see Figure 5A). The MN subfamily, previously known as Antibacterial Factors (ABFs; Tarr, 2012b), was the most widely distributed, identified in almost all clades examined (see Figure 5A). MNs are characterised by a cysteine motif consisting of eight cysteine residues which form four disulfide bonds (see Figure 4B). In line with the Diapausins, high amino acid variability exists between the key MN cysteine residues, which may contribute to diversity in antimicrobial activity. MN antimicrobial activity has only been investigated in *A. suum* ASABF-α and *C. elegans* ABF-1 and −2 which possess potent antibacterial and antifungal activity (Kato and Komatsu, 1996; Kato et al., 2002). MNs were identified in every nematode clade except for Clade 12/IV, suggesting that they may have been acquired by an ancestral nematode and subsequently lost in the genera now lacking MN-encoding genes (Clade 2/I *Trichinella* and *Trichuris* spp., Clade 8/III filarial species, and all Clade 12/IV species). The reduced distribution of the MN superfamily is consistent with the broadly reduced AMP complement in Clade 2/I and 8/III suggesting a negative selective pressure for AMP-encoding genes in these clades, whereas Clade 12/IV displays other Defensin subfamilies such that the lack of MNs could be due to functional redundancy.

**Figure 5:**
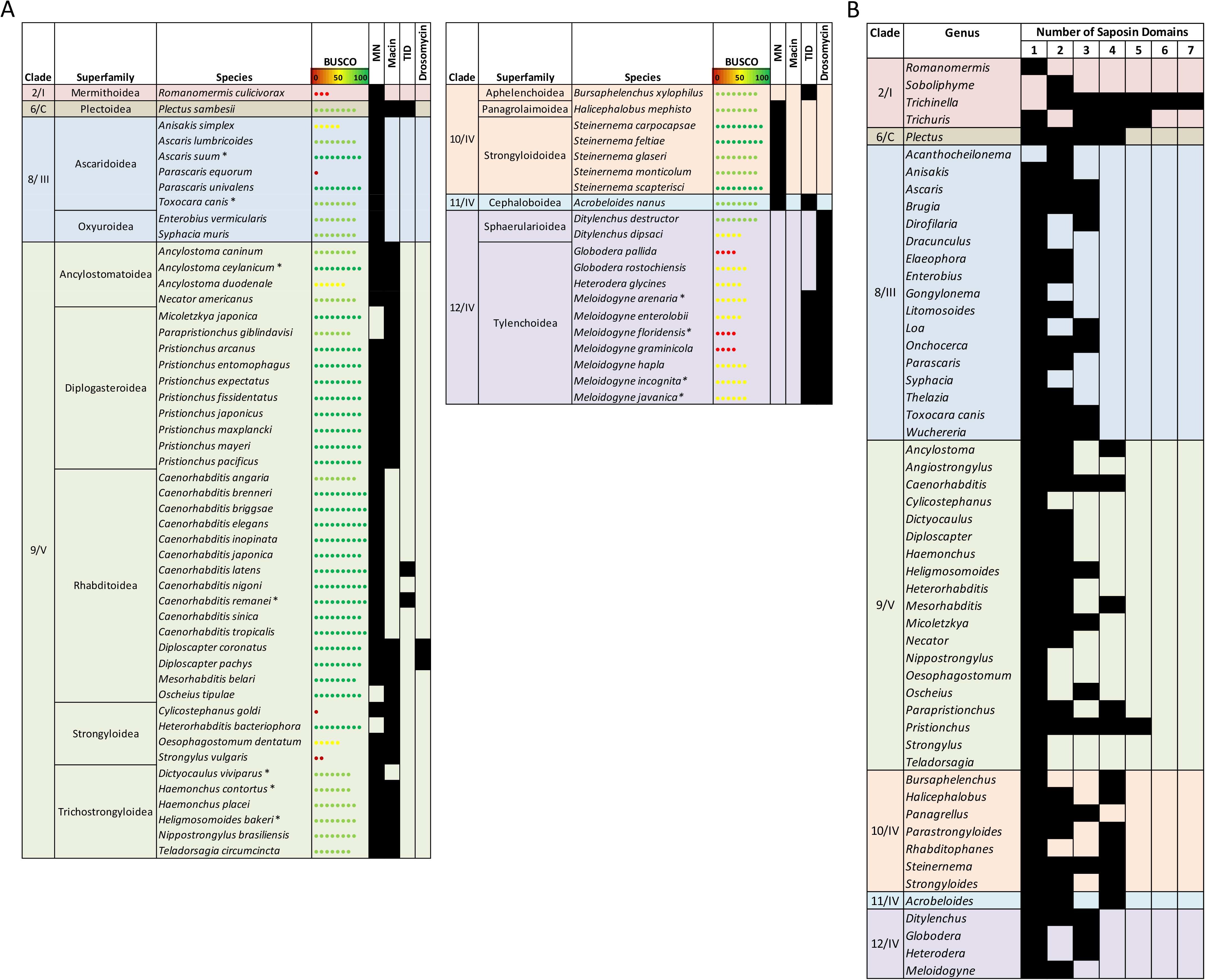
The distribution of Defensin subfamilies and multi-domain Nemapores reveal additional AMP family diversity across nematode clades. A: The distribution of four Defensin subfamilies [Mollusc/Nematode Defensin (MN), Macin, Traditional Insect Defensin (TID), Drosomycin] in Defensin-containing nematode species. B: The distribution of multi-domain Nemapore-encoding genes across nematode genera. Nematode species/genera are organised by clade (Blaxter et al., 1998, van Megan et al 2009) and superfamily designation (NCBI taxonomy browser; https://www.ncbi.nlm.nih.gov/taxonomy). Black shading indicates the presence of at least one AMP gene identified via HMM, BLASTp and tBLASTn analyses. BUSCO complete scores were obtained from Wormbase Parasite V14 (https://parasite.wormbase.org/index.html; Bolt et al 2018). Asterisk indicates multiple genome assemblies were used. Note that species which do not possess Defensin genes are not included in A.

A MN subfamily member with only six cysteine residues and a rearranged disulfide bonding pattern, known as ASABF-6Cys-α, was previously identified in *A. suum* (Minaba et al., 2009). Previous analysis of EST and draft genomic datasets identified homologs of ASABF-6Cys-α beyond Ascaridida in *Ancylostoma ceylanicum, Caenorhabditis briggsae, Caenorhabditis remanei* and *Meloidogyne hapla* (Tarr, 2012a). However, this study has revealed that the *A. ceylanicum, C. briggsae and C. remanei* sequences are, in fact, not homologs of ASABF-6Cys-α but are traditional MN Defensins which were previously incomplete. The *M. hapla* sequences were not included in this analysis as they lack key MN motifs. In this study, a single putative ortholog of *A. suum* ASABF-6Cys-α was identified in *Parascaris univalens* supporting the original hypothesis that the 6-cysteine variation in the MN Defensin sequence is only present in the Ascaridida (Minaba et al., 2009).

Nematode Macin-encoding genes have been previously reported by Gerdol et al. (2012) however this study provides the first profile of these genes across phylum Nematoda. Macin-encoding genes were identified in Clades 6/C and 9/V. Within Clade 9/V, Macin-encoding genes are found in all species except all *Caenorhabditis* spp., *Dictyocaulus viviparus* and both *Angiostrongylus* spp.. Notably, Macin-encoding genes are found in parasites of health and economic importance including *H. contortus, N. americanus* and *T. circumcincta*. The absence of Macin-encoding genes from the *Caenorhabditis* genera suggests that they have been lost in these species while they have been retained in other free-living Clade 9/V nematodes. The reason for this loss is unclear but reemphasises the need to conduct AMP research beyond *C. elegans*.

Macins from other invertebrates possess antibacterial and antifungal activities and can promote regeneration of neuronal tissue in annelids (Jung et al., 2012), however nematode Macins have not been functionally characterised. Macins typically display eight cysteine residues that form four disulfide bonds, however nematode Macins have an additional cysteine pair forming another putative disulfide bond (see Figure 4B). Theromacin from the annelid *Hirudo medicinalis*, which possesses activity against gram positive bacteria and promotes neuron regeneration, also has an additional cysteine pair suggesting that nematode Macins may possess similar functional profiles.

Drosomycin Defensins possess eight cysteine residues which form four disulfide bonds to produce a tertiary structure that is distinct from the MN Defensins (see Figure 4B). Drosomycin-encoding genes were previously identified in free-living *Panagrolaimus superbus* (Clade 10/IV) EST datasets (Tarr, 2012a). In this study, there were no *Panagrolaimus* genomes included in the nematode database however, the addition of four *Panagrolaimus* genomes in a subsequent Wormbase Parasite update (Version 15, October 2020) allowed the confirmation of Drosomycin genes in the *Panagrolaimus* genera (data not shown).

Drosomycin-encoding genes were also identified in Clade 12/IV species and two Clade 9/V species (*Diploscapter coronatus* and *Diploscapter pachys*). It is unclear whether other Clade 10/IV species have lost these genes or whether the *Panagrolaimus* species acquired them independently. Nematode Drosomycin peptides have yet to be functionally characterised however insect Drosomycin peptides display potent antifungal activity against filamentous fungi (Zhang and Zhu, 2009). Note that cysteine-rich nematode peptides have previously been designated as Drosomycin peptides but have since been reclassified as TID subfamily members based on their cysteine motif positioning (Zhu and Gao, 2014; Tarr, 2016).

Traditional Insect Defensin peptides are characterised by the typical insect Defensin cysteine array where six cysteines form three disulfide bonds (see Figure 4B). Traditional Insect Defensins display a more limited distribution across phylum Nematoda (28 TID-encoding genes in 12 nematode species). Intriguingly, the 12 nematode species that possess TID-encoding genes are phylogenetically distinct FLN and PPN species that span five nematode clades; this raises questions about the evolutionary history of these genes. The TID subfamily members appear to have functionally diversified such that each peptide has a specific activity range. For example, *Caenorhabditis remanei* Cremycin-5 possesses antifungal activity whereas Cremycin-15 lacks antifungal activity but is bioactive against gram positive bacteria (Zhu and Gao, 2014).

### 3.8. Nemapores are the most widely distributed AMP family, and can be delineated based on Saposin domain characteristics

Nemapores are cysteine-stabilised AMPs found in a wide range of animal species. They are members of the SAPosin-LIke Protein (SAPLIP) superfamily which includes other membrane-interacting proteins such as metallophosphoesterases and cell proliferator regulators (Bruhn, 2005). Nemapores are characterised by a signal peptide sequence followed by at least one Saposin domain. Saposin domains comprise of six cysteine residues in a specific arrangement (see Figure 4C). This study updates previous analyses which indicate that Nemapores are the most widely distributed AMP family in phylum Nematoda (see Figure 2; Tarr, 2012a). Indeed, at least one Nemapore-encoding gene (~15.34 ±1.38 genes per genome) was identified from every species analysed, suggesting that Nemapores are a key component of nematode innate immunity. Nemapore-encoding genes, which were previously reported as absent in *N. americanus, Strongyloides stercoralis* and *Meloidogyne incognita* (Tarr, 2012a), were identified in this study reemphasising the value in utilising updated genomic datasets.

Despite broad representation across the phylum, the numbers of Nemapore-encoding genes varies considerably across clades and species. Clades 6/C, 9/V and 10/IV appear to be enriched for Nemapore-encoding genes; for example, *Plectus sambesii* has >100 Nemapore-encoding genes. Nemapore-encoding gene expansions imply that there may be diversification of Nemapore function in these species. As discussed previously Nemapores have already been identified in the ESP of *S. feltiae* activated juveniles and this, coupled with the expansion of Nemapore-encoding genes in Clade 10/IV, highlights that Nemapores may play a key role in Clade 10/IV nematode host infection. Additionally, *C. elegans* Nemapores have been implicated in nutrition, where knockdown of *Ce-spp-5* resulted in a diminished capability to kill ingested *Escherichia coli* resulting in impeded development (Roeder et al., 2010).

There is also variation in the number of Saposin domains encoded by Nemapore genes across nematode genera (see Figure 5B). Previous analyses indicate that Nemapores containing three and four Saposin domains were restricted to species from Clades 2/I and 9/V (Tarr, 2012a). Data generated in this study, however, show that multi-domain Nemapores are more widespread where Nemapores with three or more Saposin domains are represented in all clades. Indeed, *Trichinella* spp. appear to be enriched for multi-domain Nemapores with some genes containing seven Saposin domains; multi-domain Nemapore genes have yet to be functionally characterised. In the amoeba *Naegleria fowleri* multiple bioactive Saposin peptides are processed from larger multipeptide precursor sequences (Herbst et al., 2004). It is also possible that nematode multi-domain Nemapores are post translationally cleaved to produce multiple bioactive single domain Nemapores however previous attempts to unravel the processing of a four Saposin domain protein from *Trichinella spiralis* were unable to resolve protein cleavage (Selkirk et al., 2004).

### 3.9. Expansion of the GRSPs is not restricted to the Caenorhabditis genera

This study provides the first pan-phylum profile of the GRSP-encoding genes across phylum Nematoda; previously GRSPs had only been profiled in *C. elegans* and *C. briggsae* where they were found to be enriched relative to several other model organisms (Ying et al., 2016). The data presented here demonstrates that GRSPs are, in fact, the most abundant AMP family in phylum Nematoda (3145 genes; 23.47 ±2.26 genes per genome) with Clades 9/V and 10/IV exhibiting enriched profiles. This evolutionary expansion of GRSPs extends beyond *Caenorhabditis* genera (see Figure 2), where GRSP-encoding genes are absent from only two nematode species [*Dracunculus medinensis* and *Litomosoides sigmodontis* (Clade 8/III)] which suggests importance to nematode biology. Functional characterisation of GRSPs has been restricted to *C. elegans* where their expression has been profiled post microbial infection (Ying et al., 2016), and antibacterial and antifungal assays reveal antimicrobial properties that do not involve membrane permeabilisation (Couillault et al., 2004; Lim et al., 2016). Further work will reveal if this characteristic is conserved across nematode GRSPs. In addition, it is interesting to note that a number of Neuropeptide-Like Proteins (NLPs) have been classified as GRSPs (NLP-10, NLP-12, NLP-21 and NLP-24-34), highlighting a need to unravel the role of these peptides.

### 3.10. Stage specific transcriptome data highlight that AMPs are expressed in key nematode life stages

To assess the biological relevance of AMPs across nematode life stages, publicly available life stage specific RNAseq datasets were examined for 15 species (10 APN, 1 EPN, 3 PPN, 1 FLN). >85% of AMP genes were expressed (TPM >2) in at least one life stage (see Figure 1E). Notably, despite significant variation in the total number of AMP genes encoded within individual nematode species, this did not translate to major variability in the proportion of total AMP genes expressed. For example, in *Trichuris muris* six of the seven (86%) identified AMP genes were expressed whereas in *P. pacificus* 88 of the 102 (86%) identified AMP genes were expressed. There appear to be differences in the expression of AMP genes from the same AMP family such that no AMP family is consistently expressed in a specific life stage in the five parasitic species examined in more detail (see Supplementary Figure S1). This suggests that closely related AMPs may have functionally diversified and reinforces the need to further characterise these families in other species. Notably, in *A. suum* and *H. contortus* there is clear elevation in the expression of AMP genes in life stages which are more exposed to the host gastrointestinal system further supporting that these genes may play a role in nematode innate immunity.

### 3.11. Improved genome and transcriptome data will drive AMP discovery and functional characterisation in phylum Nematoda

Genome and transcriptome studies possess inherent caveats that make extrapolation of the data challenging. For example, in this study there are significant differences in the quality of the genome datasets where some chromosomal level assemblies (e.g. *Strongyloides ratti*) are available while others remain at contig level (e.g. *M. hapla*). The quality of genome datasets was considered through the integration of Benchmarking Universal Single-Copy Ortholog (BUSCO) complete scores for each genome (see Figure 2; Simão et al., 2015) such that there appears to be a correlation between the number of AMPs genes identified and genome quality as denoted by BUSCO score; for example, 7 AMP genes in *Parascaris equorum* (BUSCO complete score of 5) relative to 31 AMP genes in the closely related *Parascaris univalens* (BUSCO complete score of 90.9). As nematode genome assemblies and annotations are improved, more AMP genes will be uncovered to reveal a more comprehensive nematode antimicrobial peptidome.

The homology-derived approach used in this study to identify AMPs was driven by query sequences primarily derived from *C. elegans* and *A. suum*, such that the HMM models are likely to be more biased towards the identification of *Caenorhabditis*- or *Ascaris*-like AMPs. Consequently, it is possible that AMPs which have evolved rapidly in distantly related species may be overlooked using this approach, underscoring the need to develop alternative strategies for *in silico* AMP discovery. Indeed, it is difficult to assess the significance of the apparent reduced AMP profiles in Clades 2/I, 8/III and 12/IV until comprehensive AMP discovery approaches have been deployed across phylum Nematoda. Verification of AMPs in nematode tissues and biofluids using peptidomics technologies (Atkinson et al., 2021) will seed follow-on functional biology.

## 4. Conclusions

This study provides a comprehensive genome-level pan-phylum profile of nematode AMP-encoding genes which will inform fundamental nematode biology and seed novel antimicrobial discovery. The data demonstrate that nematodes have a rich AMP complement that includes 5887 AMP-encoding genes in 109 species, ~90% of which are novel. The pan-phylum approach has revealed that APNs and PPNs possess fewer AMP-encoding genes than FLNs suggesting that parasitic nematodes may possess a more specific AMP arsenal tailored towards the host environments they inhabit. Major differences in the distribution and abundance of AMP families were evident across the phylum where Clade 2/I, 8/III and 12/IV species possess fewer AMP-encoding genes, while Clade 9/V and 10/IV species have expanded AMP repertoires. Two AMP families, Cecropins and Diapausins, appear to be restricted to Ascarids and FLNs respectively whereas Defensins, Nemapores and GRSPs were more widely distributed across the phylum. This study provides a springboard for functional biology to unravel the importance of AMPs to parasite and infection biology and will inform future drug discovery pipelines.

## Acknowledgments

The authors acknowledge support for this work from: Biotechnology and Biological Sciences Research Council grant (BB/H019472/1), Biotechnology and Biological Sciences Research Council/Boehringer Ingelheim grants (BB/MO10392/1, BB/T016396/1), and Department of Agriculture Environment and Rural Affairs Northern Ireland (DAERA).

## Supplementary Data

**Supplementary Table S1: AMP query sequences used for construction of AMP family HMMs.**

**Supplementary Table S2: Genomic and transcriptomic datasets used in the study.**

**Supplementary Table S3: AMP gene IDs identified from 134 nematode predicted protein datasets.**

**Supplementary Table S4: Number and relative abundance of AMP genes identified in this study.**

**Supplementary Figure S1: Summary of transcriptomic data.** Life stage specific heatmaps of AMP expression for AMP genes from B: *Trichuris muris* (6 genes), C: *Ascaris suum* (32 genes), D: *Haemonchus contortus* (58 genes), E: *Strongyloides stercoralis* (22 genes), F: *Meloidogyne incognita* (36 genes). Heatmaps generated using Heatmapper (http://www.heatmapper.ca/; Babicki et al., 2016). Expression is shown in Z scores and scaled by row. Genes are clustered using Average Linkage and Euclidean distance method. Darker shades represent higher expression and lighter shades represent lower expression.

