## Supplementary Figure S1 for "The Nematode Antimicrobial Peptidome: a novel opportunity for parasite control?"

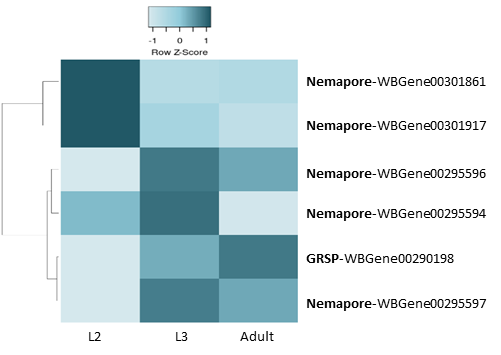

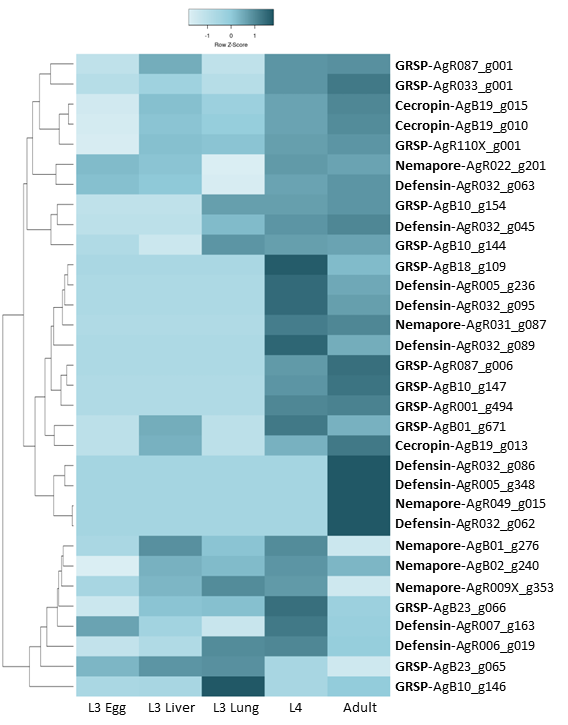

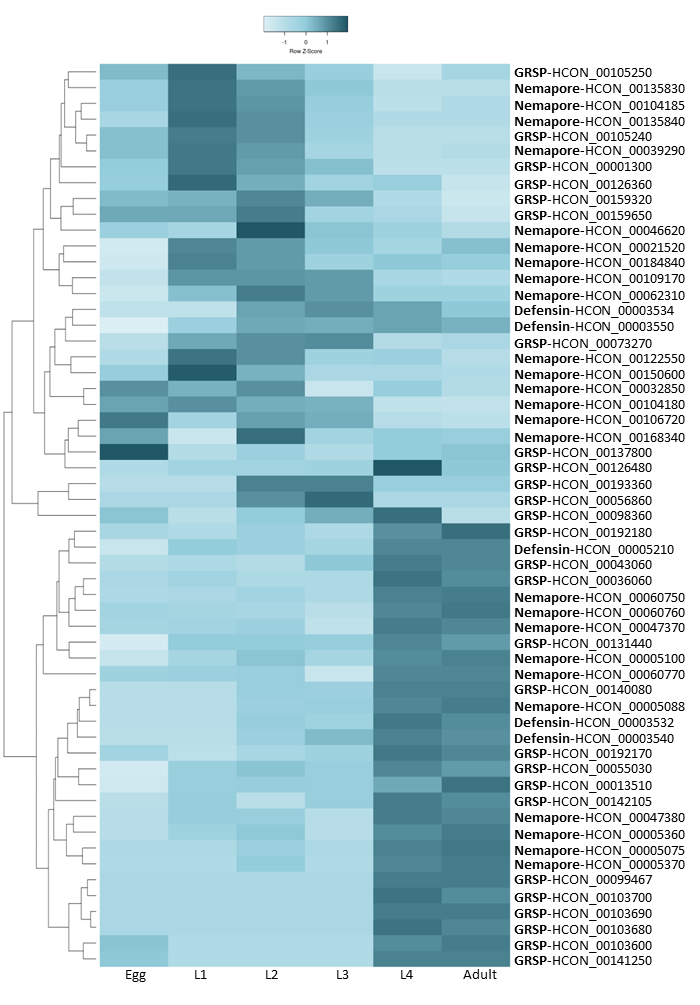

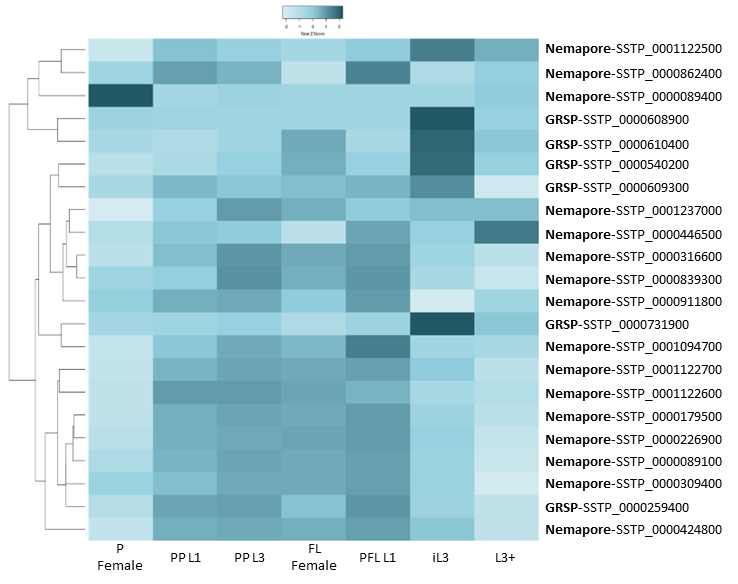

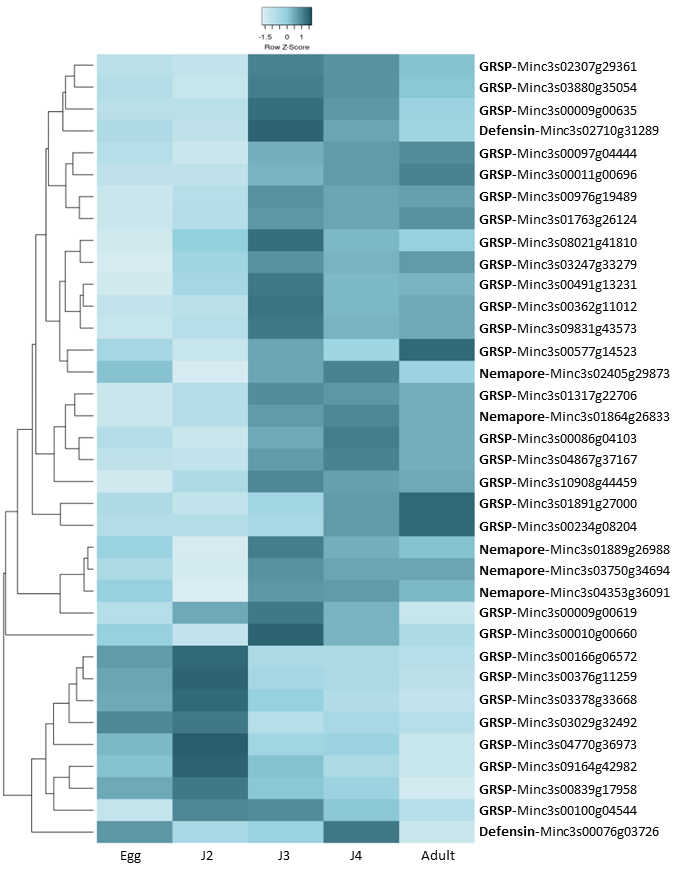

A. *Trichuris muris*

B. *Ascaris suum*

C. *Haemonchus contortus*

D. *Strongyloides stercoralis*

E. *Meloidogyne incognita*
